# Cell-type specific epigenetic links to schizophrenia risk in brain

**DOI:** 10.1101/609131

**Authors:** Isabel Mendizabal, Stefano Berto, Noriyoshi Usui, Kazuya Toriumi, Paramita Chatterjee, Connor Douglas, Iksoo Huh, Hyeonsoo Jeong, Thomas Layman, Carol Tamminga, Todd M. Preuss, Genevieve Konopka, Soojin V. Yi

**Author notes:** Co-first author (Isabel Mendizabal, Stefano Berto, Noriyoshi Usui, Kazuya Toriumi). Correspondence (G.K.), (S.V.Y.).

## Abstract

The importance of cell-type specific epigenetic variation of non-coding regions in neuropsychiatric disorders is increasingly appreciated, yet data from disease brains are conspicuously lacking. We generated cell-type specific whole-genome methylomes (*N*=95) and transcriptomes (*N*=89) from neurons and oligodendrocytes from brains of schizophrenia and matched controls. The methylomes of these two cell-types are highly distinct, with the majority of differential DNA methylation occurring in non-coding regions. DNA methylation difference between control and schizophrenia brains is subtle compared to cell-type difference, yet robust against permuted data and validated in targeted deep-sequencing analyses. Differential DNA methylation between control and schizophrenia tends to occur in cell-type differentially methylated sites, highlighting the significance of cell-type specific epigenetic dysregulation in a complex neuropsychiatric disorder. Our resource provides novel and comprehensive methylome and transcriptome data from distinct cell populations from schizophrenia brains, further revealing reduced cell-type epigenetic distinction in schizophrenia.

## Introduction

Schizophrenia is a lifelong neuropsychiatric psychotic disorder affecting 1% of the world’s population^1^. Genetic dissection of schizophrenia risk has revealed the polygenic nature of the disorder^2–4^. Many of the schizophrenia risk loci are located in non-coding regions of the genome, suggesting gene regulation plays a role in disease pathology. Indeed, a large number of these risk loci are associated with alterations in gene expression in schizophrenia^2, 5, 6^. These observations implicate epigenetic mechanisms as potential mediators of genetic risk in schizophrenia pathophysiology. Epigenetic mechanisms, such as DNA methylation, may have particular relevance for human brain development and neuropsychiatric diseases^7–9^. Previous studies found that changes in DNA methylation associated with schizophrenia are significantly enriched with DNA methylation changes from prenatal to postnatal life^7^. Moreover, genome-wide association studies (GWAS) of schizophrenia risk loci were over-represented in variants that affect DNA methylation in fetal brains^10^.

Prior studies of the genetic and epigenetic risks for schizophrenia have some limitations, however, including the use of pre-defined microarrays, which traditionally focused on CpG islands and promoters, for methylation profiling. Unbiased, genome-wide analyses of DNA methylation are revealing that variation in DNA methylation outside of promoters and CpG islands define critical epigenetic difference between diverse cell-types^11, 12^. Additionally, previous genomic studies of schizophrenia have used brain tissue samples containing a heterogeneous mixture of cell-types, although gene expression patterns vary considerably across cell-types in the human brain^13–17^. To address these concerns, we carried out whole-genome methylome and transcriptome analyses using post-mortem human brain tissue that underwent fluorescence-activated nuclei sorting (FANS)^18^ into neuronal (NeuN^+^) and oligodendrocyte (OLIG2^+^) cell populations. Both neurons and myelin-forming oligodendrocytes have been implicated in schizophrenia pathophysiology^19, 20^, and may be functionally dependent on one another for proper signaling in the brain^21^. Tissue was dissected from Brodmann area 46 (BA46) of the dorsolateral prefrontal cortex, a key brain region at risk in schizophrenia^1, 22^. We used whole-genome bisulfite sequencing (WGBS) to obtain an unbiased assessment of epigenetic modifications associated with schizophrenia, and additionally carried out whole-genome sequencing (WGS) and RNA-sequencing (RNA-seq) of the same samples to document transcriptomic consequences while accounting for genetic background differences.

Integrating these data, we demonstrate extensive differential DNA methylation between neurons and oligodendrocytes. Comparisons to previous studies using bulk tissues indicate that they were generally biased toward neuronal populations. Our resource thus offers comprehensive and balanced analyses of molecular variation in control and disease brains, including novel information from a major yet relatively underexplored brain cell population (oligodendrocytes). This comprehensive and novel data set allows us to demonstrate subtle yet robust DNA methylation differences between control and schizophrenia samples, which are highly enriched in sites that are epigenetically differentiated between the two cell-types. Moreover, we show that schizophrenia associated DNA methylation changes reduce the cell-type methylation difference. Together, these data indicate that the integration of multiple levels of data in a cell-type specific manner can provide novel insights into complex genetic disorders such as schizophrenia.

## Results

### Divergent patterns of DNA methylation in human brain cell-types

We performed FANS^18^ using post-mortem tissue dissected from BA46 of the dorsolateral prefrontal cortex using NeuN and OLIG2 antibodies (Fig. 1a; Methods). Immunofluorescent labeling indicates that NeuN-positive nuclei and OLIG2-positive nuclei following FANS (hereinafter “NeuN^+^” or “OLIG2^+^”) represent neurons within the cerebral cortex and oligodendrocytes and their precursors, respectively (Figure 1b,c,d). We analyzed genomic DNA (gDNA) and total RNA from the same nuclei preparations in NeuN^+^ or OLIG2^+^ by WGBS and RNA-seq. We additionally carried out WGS of the brain samples to explicitly account for the effect of genetic background differences.

**Figure 1.**
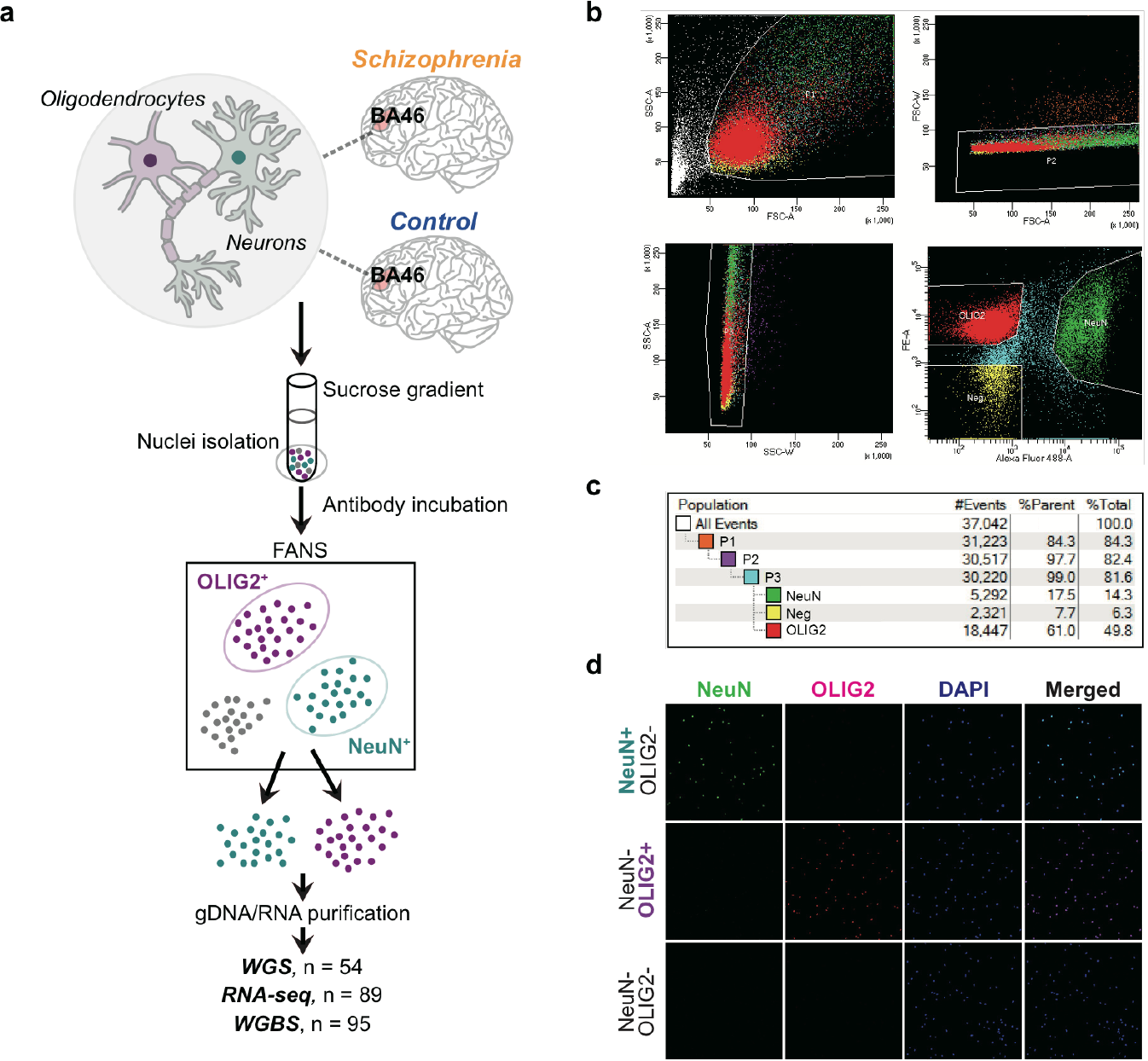
Experimental design and FANS workflow example. **a.** Post-mortem brain tissue from BA46 was matched between cases with schizophrenia and unaffected individuals. Tissue pieces were processed to isolate nuclei and incubated with antibodies directed towards NeuN or OLIG2. The nuclei were sorted using fluorescence-activated nuclei sorting (FANS) to obtain purified populations of cell-types. Nuclei were processed to obtain genomic DNA (gDNA) and nuclear RNA from the same pools. Nucleic acids then underwent whole-genome sequencing (WGS), whole-genome bisulfite sequencing (WGBS), or RNA-sequencing (RNA-seq). **b.** NeuN-positive (NeuN^+^) nuclei represent neurons within the cerebral cortex as few human NeuN-negative (NeuN^−^) cells in the cortex are neurons^30, 31^ (e.g. Cajal-Retzius neurons). OLIG2-positive (OLIG2^+^) nuclei represent oligodendrocytes and their precursors^32, 33^. Isolation of nuclei expressing either NeuN conjugated to Alexa 488 or OLIG2 conjugated to Alexa 555. Nuclei were first sorted for size and complexity, followed by gating to exclude doublets that indicate aggregates of nuclei, and then further sorted to isolate nuclei based on fluorescence. “Neg” nuclei are those that are neither NeuN^+^ nor OLIG2^+^. **c.** Example percentage nuclei at each selection step during FANS. Note that while in this example more nuclei were OLIG2^+^, in other samples, the proportions might be reversed. **d.** Immunocytochemistry of nuclei post-sorting. Nuclei express either NeuN or OLIG2 or are negative for both after FANS. DAPI labels all nuclei.

Whole-genome DNA methylation maps of NeuN^+^ (*N*=25) and OLIG2^+^ (*N*=20) from control individuals (Supplementary Table 1) show a clear separation of the two populations (Fig. 2a). Previously published whole-genome methylation maps of neurons^23^ co-segregate with NeuN^+^. On the other hand, previously generated NeuN^−^-methylomes^23^ cluster as outliers of OLIG2^+^ samples, potentially due to the inclusion of other cell-types compared to our cell-sorted samples. We identified differentially methylated CpGs between cell types, which we refer to as ‘differentially methylated positions (DMPs)’, using a statistical method that allows us to explicitly take into account the effect of covariates (Supplementary Table 2, Methods), while handling variance across biological replicates as well as the beta-binomial nature of read count distribution from WGBS^24^. Despite the large number of CpGs (~26 million total), we identify numerous DMPs between NeuN^+^ and OLIG2^+^ after correcting for multiple testing. At a conservative Bonferroni *P* < 0.05, over 4 million CpGs are differentially methylated between these two cell types, revealing highly distinct cell type difference in gDNA methylation (Fig. 2a,b). On average, DMPs between NeuN^+^ and OLIG2^+^ exhibit a 32.6% methylation difference. NeuN^+^ tends to be more hyper-methylated than OLIG2^+^ (Fig. 2b; 64% of DMPs, binomial test, *P*< 10^−16^). This observation is consistent with NeuN^+^ being more hyper-methylated than non-neuronal populations^23^.

**Figure 2.**
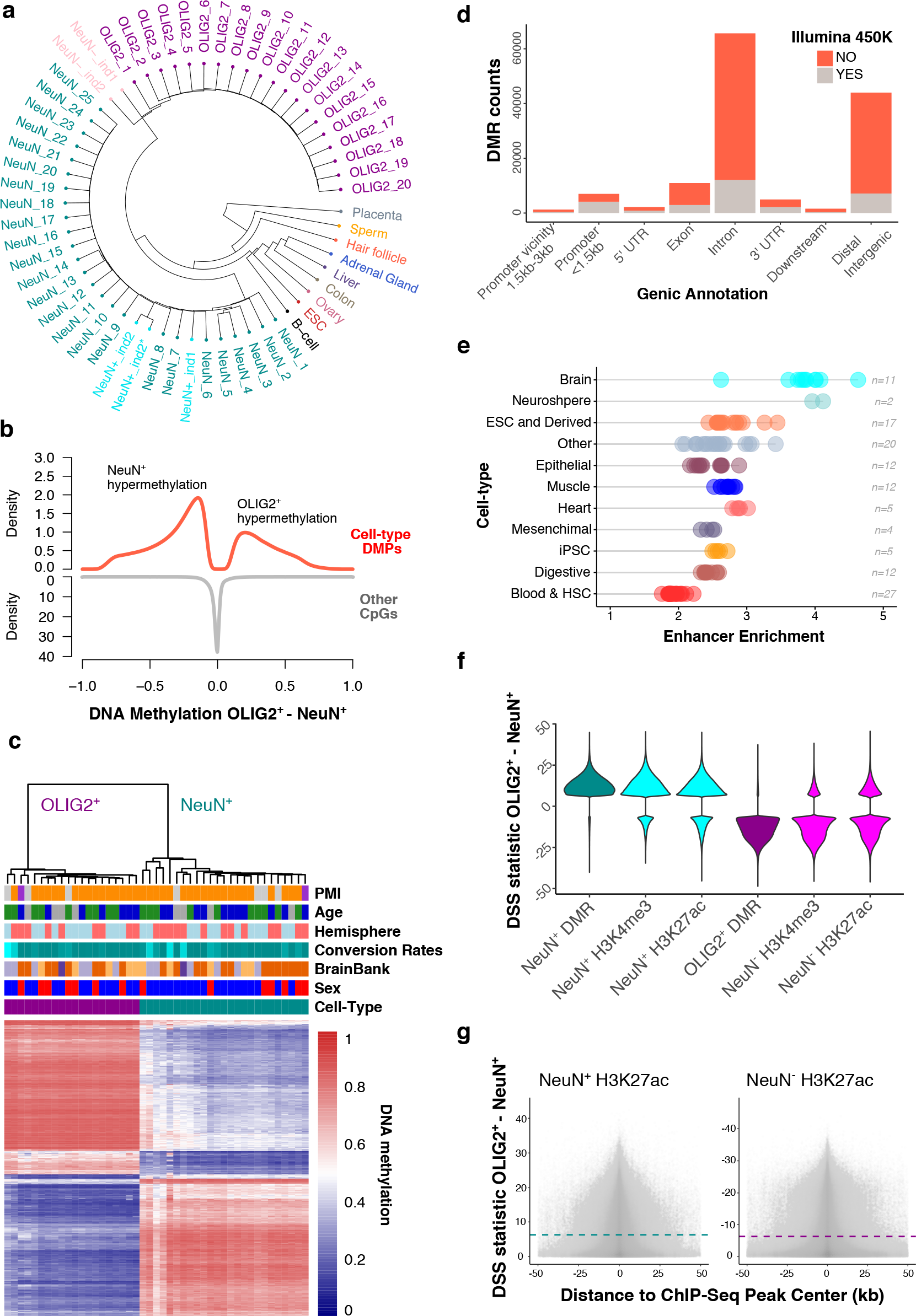
Divergent patterns of DNA methylation in NeuN^+^ and OLIG2^+^ cell-types in the human brain. **a.** Clustering analysis based on whole-genome CpG methylation values completely discriminated between NeuN^+^ (*N*=25) and OLIG2^+^ (*N*=20) methylomes. Additional NeuN^+^ (colored in turquoise) and those labeled as NeuN^−^ (pink) are from^23^. **b.** Density plots showing the distribution of fractional methylation differences between OLIG2^+^ and NeuN^+^ at differentially methylated positions (DMPs) and other CpGs (non-DMPs). We observed a significant excess of NeuN^+^ hypermethylated DMPs compared to OLIG2^+^ (binomial test with expected probability = 0.5, *P* < 10^15^). **c.** Heatmap of top 1,000 most significant DMRs between OLIG2^+^ and NeuN^+^. Fractional methylation values per individual (column) and DMR (row) show substantial differences in DNA methylation and clear cell-type clustering. **d.** Genic annotation of DMRs and coverage with Illumina 450K Methylation Arrays. Counts of different genic positions of DMRs are shown. DMRs containing at least one CpG covered by a probe in the array are indicated. Only 20.8% of the DMRs contain one or more CpG targeted by Illumina 450K array probes. **e.** DMRs are enriched for brain enhancers. Enrichment of enhancer states at DMRs compared to 100 matched control DMR sets from 127 tissues^28^. Random sets are regions with similar characteristics as, including the total number of regions, length, chromosome and CG content. **f.** Correspondence between cell-type specific methylation sites in NeuN^+^ and OLIG2^+^ with NeuN^+^ and NeuN^−^ ChIP-seq data sets^9^. Neuron-specific ChIP-seq peaks show an excess of sites with NeuN^+^ specific hypomethylated sites (positive DSS statistic, see Methods) whereas non-neuron peaks showed significant enrichment for sites specifically hypomethylated in OLIG2^+^ (negative DSS statistic). **g.** Distribution of cell-type specific methylation differences at CpGs within H3K27ac ChIP-seq peaks in NeuN^+^ and NeuN^−^ nuclei. Positive values of DSS statistic indicate hypomethylation in NeuN^+^ compared to OLIG2^+^, whereas negative values indicate hypermethylation (see Methods). Dashed lines indicate the significance level for DSS analyses.

As expected from regional correlation of DNA methylation between adjacent sites^25–27^, many DMPs occur near each other, allowing us to identify ‘differentially methylated regions’ or ‘DMRs’ (defined as ≥5 significant DMPs in a 50 bp region) spanning 103 MB in the human genome, exhibiting mean methylation difference of 38.3% between cell types (Figure 2c, Supplementary Table 3). Many DMRs reside in introns and distal intergenic regions (Figure 2d), which are traditionally viewed as ‘non-coding’. Chromatin state maps based on six chromatin marks^28^ indicate that many DMRs, especially those located in introns and distal intergenic regions, exhibit enhancer chromatin marks, in particular, brain enhancers (OR between 2.6 and 4.6 fold, *P* < 0.01, Fig. 2e and Supplementary Table 4). In fact, over 60% of all DMRs show enhancer-like chromatin features in brain (Supplementary Fig. 1). These results highlight the regulatory significance of non-coding regions of the genome. Notably, currently available arrays such as the Illumina 450K do poorly in terms of targeting putative epigenetic regulatory loci (Fig. 2d).

NeuN^+^-specific hypo-methylated regions are significantly enriched in recently identified NeuN^+^ specific H3K4me3 and H3K27ac peaks^9^ (Fig. 2f; Fisher’s exact test OR = 7.8, *P* < 10^−15^). H3K4me3 and H3K27ac peaks in the NeuN^−^ populations also show significant enrichment of OLIG2^+^-specific hypo-methylation, although the degree of enrichment is less strong than NeuN^+^ correspondence (Fisher’s exact test OR = 4.8, *P* < 10^−15^), again potentially due to the inclusion of other types of cells. WGBS data is complementary to ChIP-seq data in terms of resolution and coverage. While ChIP-seq provides resolution in the scale of several thousand base pairs (for example, peak sizes in previous study^9^ are on average several kilobases and extend up to several hundred kilobases), WGBS data offers base-pair resolution. Even though DMPs are generally concentrated around the center of ChIP-seq peaks, some peaks show more diffuse patterns, indicating that incorporating DMP information could offer fine-scale resolution of histone modification in individual genomic regions (Fig. 2g and Supplementary Fig. 2).

We further examined DNA methylation of cytosines that are not in the CpG context, as nucleotide-resolution whole-genome DNA methylation maps have begun to reveal the potential importance of non-CG methylation (CH methylation, where H = A, C, or T) particularly in neuronal function^23^. We observed that low levels of CH methylation were present in NeuN^+^ but nearly absent in OLIG2^+^ (Supplementary Fig. 3), consistent with previous reports^23^. CH methylation is primarily associated with CA nucleotides (69.4%), followed by CT (26%), and CC (4.6%) (Supplementary Fig. 3). In addition, gene body mCH values negatively correlate with gene expression in NeuN^+^ (Spearman’s rho −0.16, *P* < 10^−10^; Supplementary Fig. 3). Therefore, CH patterns at gene bodies provide an additional layer of gene expression regulation that is specific to neurons while absent in oligodendrocytes in the human brain.

### Strong association between cell-type specific DNA methylation and expression

We next performed RNA-seq using RNAs extracted from the nuclei-sorted populations (Methods). NeuN^+^ and OLIG2^+^ transcriptomes form distinctive clusters (Fig. 3a). Transcriptomic data from cell-sorted populations clustered closer to bulk RNA-seq data from cortical regions but were distinct from those from cerebellum and whole blood (Supplementary Fig. 4). We further show that previously generated bulk RNA-seq data^5, 6^ have higher proportion of NeuN^+^ compared with OLIG2^+^ (Fig. 3b), indicating that these previously generated bulk RNA-seq data are biased toward neurons. The higher neuronal proportion in bulk RNA-seq is highlighted also using an independent single-nuclei data (Supplementary Fig. 5).

**Figure 3.**
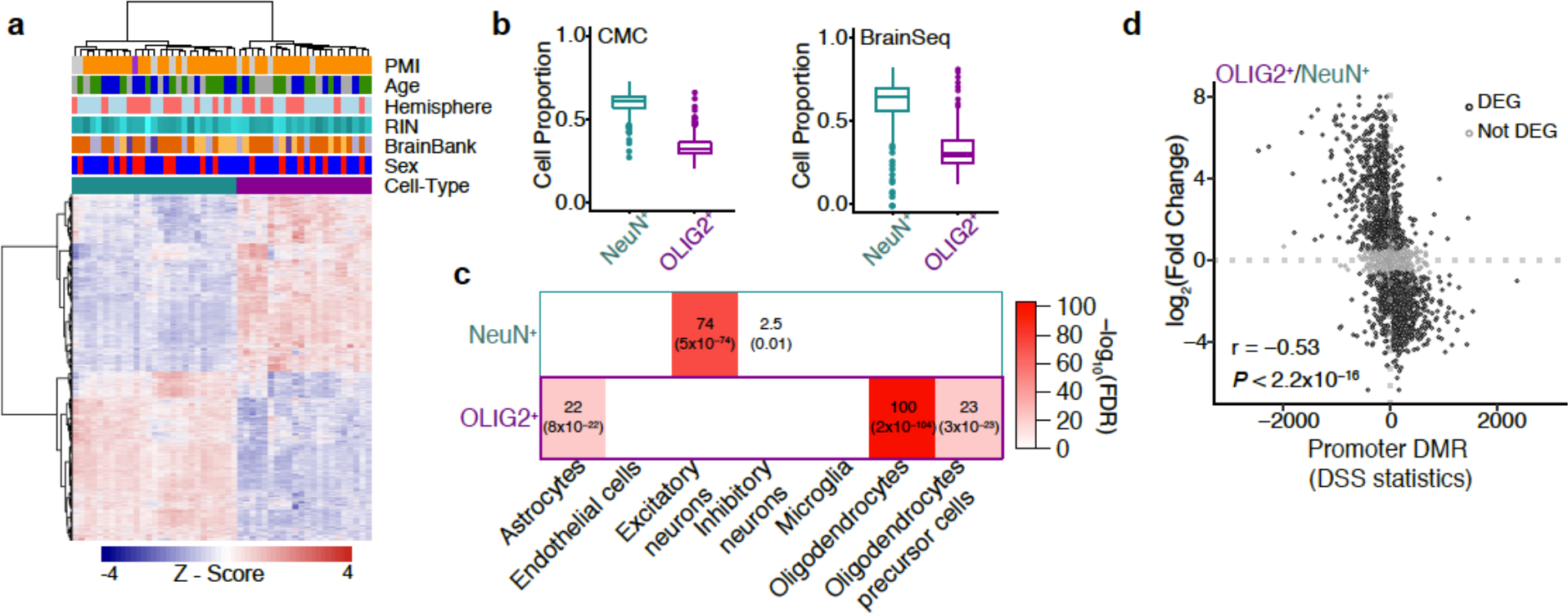
Gene expression signatures in NeuN^+^ and OLIG2^+^ nuclei. **a.** Heatmap of cell-type DEGs with covariates indicated. **b**. Cell deconvolution of bulk RNA-seq data from the CommonMind Consortium and BrainSeq compared with NeuN^+^ and OLIG2^+^ (control samples). Y-axes show the weighed proportion of cells that explain the expression of bulk RNA-seq **c**. Gene set enrichment for cell-type markers from single-nuclei RNA-seq. Enrichment analyses were performed using a Fisher’s Exact Test. Odds ratios and FDRs (within parentheses) are shown. **d.** Correspondence between expression change and methylation change in cell-types. The x-axis represents differential DNA methylation statistics for genes harboring DMRs in promoters. The y-axis indicates the log_2_(Fold Change) of expression between the two cell types. The negative correlation supports the well-established impact of promoter hypomethylation on upregulation of gene expression.

We show that 55% of genes shows significant change in expression between NeuN^+^ and OLIG2^+^ (|log_2_(Fold Change)| > 0.5 and Bonferroni correction < 0.05; Supplementary Table 5). NeuN^+^ and OLIG2^+^ specific genes (defined as significantly up-regulated in NeuN^+^ compared to OLIG2^+^ and vice versa) are enriched for known markers of specific cell-types of brain. Specifically, NeuN^+^ specific genes are enriched for excitatory and inhibitory neurons, whereas OLIG2^+^-specific genes show strong enrichment for oligodendrocytes and lower enrichment for oligodendrocyte precursor cells and astrocytes (Fig. 3c). Divergent DNA methylation between cell-types can explain a large amount of gene expression variation between cell-types (Fig. 3d, Spearman’s rho = −0.53, *P* < 10^−15^). Significant correlation extends beyond the promoter regions (Supplementary Fig. 6),

### Differential DNA methylation associated with schizophrenia

We additionally analyzed the whole-genome methylation maps of 28 NeuN^+^ and 22 OLIG2^+^ from schizophrenia brains (Methods). Compared to the robust signal of cell-type difference, DNA methylation changes associated with schizophrenia are subtler. We find no significantly differentially methylated individual CpGs between control and schizophrenia when applying stringent multiple testing correction of Bonferroni *P* < 0.05. At a moderately stringent FDR < 0.2, we identify 261 individual CpGs (60 in NeuN^+^ and 201 in OLIG2^+^) that are differentially methylated between control and schizophrenia (referred to as ‘szDMPs’ hereinafter). Applying additional filtering for high coverage sites (20× in at least 80% of samples per disease-control group), we identify a total of 97 szDMPs (14 NeuN^+^ and 83 OLIG2^+^ specific, respectively) at FDR < 0.2 (Supplementary Tables 6-7). The majority of the schizophrenia-associated loci are intronic (50.5%), and distal intergenic CpGs (45.4%) whereas only two szDMPs (2%) are located within 3kb from transcriptional start sites (Supplementary Tables 6-7).

We assessed the robustness of the results as well as the effects of covariates or potential hidden structures in the data by permutation analysis, by randomly assigning case/control labels 100 times per cell-type. The observed DNA methylation difference between control and schizophrenia samples is significantly greater than those observed in the permuted samples (Supplementary Fig. 7). Even though our statistical cutoff is moderate, considering that we are correcting for an extremely large number of independent tests, the results from permutation analyses provide support that these sites represent schizophrenia-associated signals of differential DNA methylation. Indeed, quantile-quantile plots suggest that our data exhibit a modest but significant excess of good *P* values (Fig. 4a). We also performed targeted experiments of 66 CpGs (16 szDMPs at FDR < 0.2 and 50 adjacent sites) through deep coverage sequencing using 24 samples from the discovery cohort as well as an additional 20 new independent samples. This validation analysis achieved an average read depth coverage > 14,500X. Technical replicates are highly correlated with the fractional methylation values obtained from the WGBS (Spearman’s rho = 0.96, *P* < 10^−15^, Supplementary Fig. 8), indicating the reliability of the fractional methylation estimates obtained in the discovery WGBS data. In addition, the WGBS data and validation data are highly consistent for case-control comparisons in both sign direction and correlation in effect size (Spearman’s rho = 0.87, *P* < 10^−16^ and 81.25% sign concordance, Supplementary Fig. 9). These results support the validity of szDMPs discovered in our study.

The average methylation difference between schizophrenia versus controls at szDMPs is ~6% (Supplementary Tables 6-7). There is no direct overlap between these DMPs and those previously identified from a microarray study^7^. However, despite the lack of direct overlap, the direction of methylation change between control and schizophrenia between the two studies are largely consistent in the NeuN^+^, especially with increasing significance (decreasing *P* values) (Fig. 4b). This pattern is highly significant compared to the permuted data (Fig. 4b). In comparison, the OLIG2^+^ data set does not exhibit such a pattern (Fig. 4b), potentially because the bulk tissue samples consisted largely of neurons. Deconvolution analyses of transcriptomes using our cell-sorted population support this idea (Fig. 3b).

**Figure 4.**
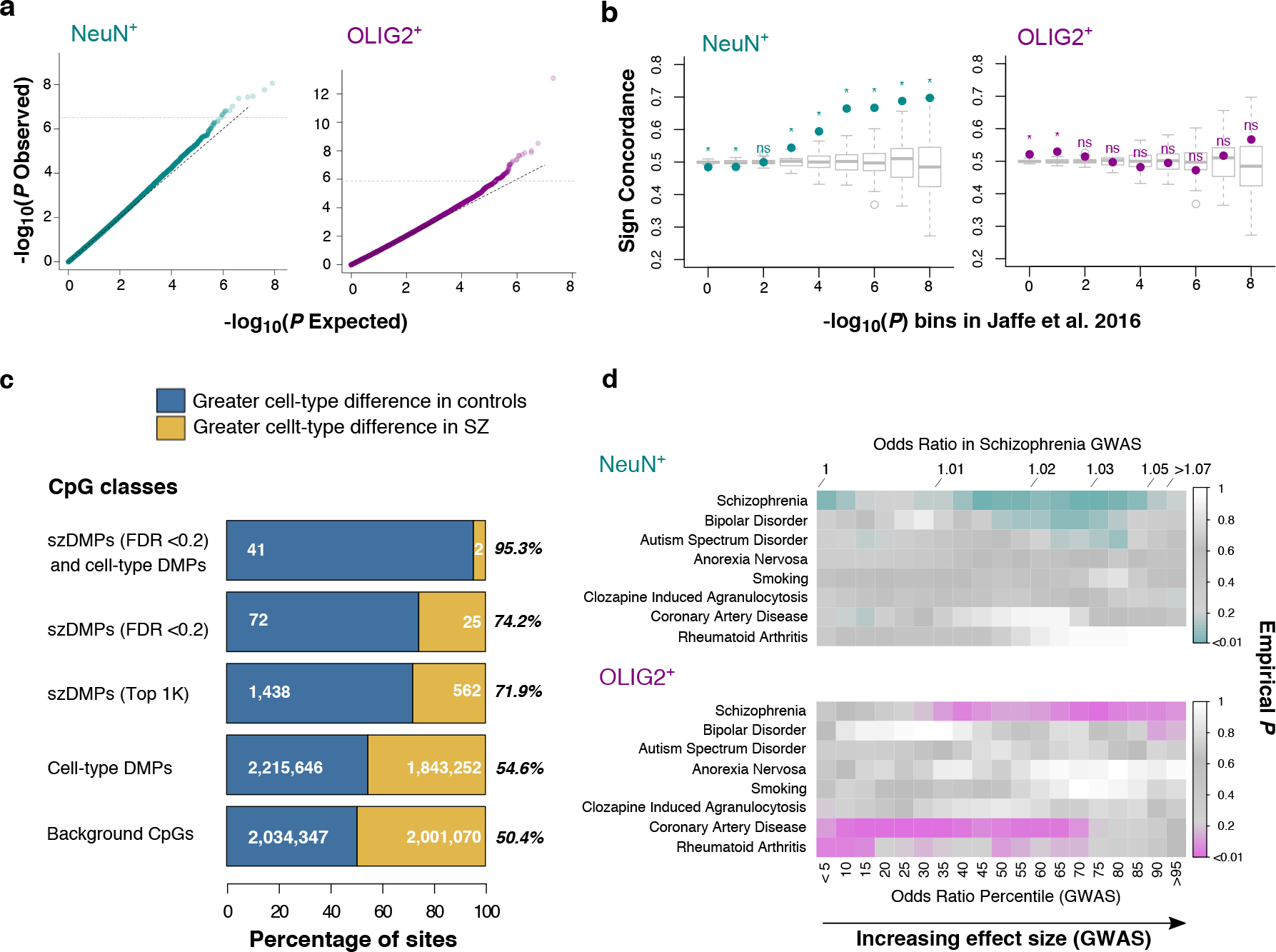
Cell-type DNA methylation patterns associated with schizophrenia. **a.** DMPs associated with schizophrenia. Quantile-quantile plots of genome-wide *P* values for differential methylation between schizophrenia and control based on NeuN^+^ (left) and OLIG2^+^ (right) WGBS data**. b.** Concordance between WGBS data and microarray-based data. Y-axis shows the ratio of sites showing concordant direction in schizophrenia versus control in our study at each *P* value bin compared with the Jaffe et al study^7^ (x-axis). Concordance was tested using a binomial test (stars indicate *P* < 0.05). Boxplots correspond to directional concordance in 100 sets of association results after case-control label permutations. NeuN^+^ (left) and OLIG2^+^ (right). **c.** szDMPs show altered cell-type differences. Barplot shows the percentage of sites with larger cell-type differences in controls than in schizophrenia and vice versa at different CpG classes. Absolute OLIG2^+^ vs NeuN^+^ methylation differences are larger in controls than cases in szDMPs compared to cell-type DMPs and non-DMP or background CpGs. szDMPs were detected as differentially methylated between cases and controls at FDR < 0.2 in NeuN^+^ (14 sites) and OLIG2^+^ samples (83 sites). Top 1K szDMPs were selected as the top 1,000 loci according to best *P* values in each cell-type (*N* = 2,000). Cell-type DMPs were detected by comparing OLIG2^+^ versus NeuN^+^ methylomes at Bonferroni *P* < 0.05. Background CpGs were sampled from CpGs showing non-significant *P* values for both case-control and OLIG2^+^ vs. NeuN^+^ comparisons. Stars represent *P* values for binomial tests with all comparisons showing *P* < 10^−7^. **d.** szDMPs are enriched for SZ GWAS signals. szDMPs identified in our methylation study in both cell-types consistently co-localize with genetic variants with moderate to large effect sizes for schizophrenia risk than expected. The table shows the empirical *P* values of szDMPs at each odds-ratio (OR) percentile of different traits from genome-wide association studies (GWAS). The actual ORs corresponding to the schizophrenia percentiles are indicated at the top.

### Enrichment of szDMPs in cell-type distinct sites imply cell-type dysregulation

Remarkably, szDMPs are significantly enriched in cell-type specific DMPs (OR = 4.1, *P* < 10^−10^, Fisher’s exact test). Similar overrepresentations are obtained when the top 1,000 szDMPs per cell-type are considered (ordered by lowest *P* value, OR = 4.05, *P* < 10^−15^, Fisher’s exact test, Supplementary Fig. 10), indicating that the enrichment is not due to the small number of szDMPs.

Moreover, szDMPs show distinct directionality in the distinct brain cell-types. Specifically, NeuN^+^ szDMPs show an excess of hypomethylation in schizophrenia samples compared to control samples (93%, 13 out of 14, *P* = 0.0018 by binomial test, Supplementary Fig. 7). An opposite pattern is observed for OLIG2^+^ szDMPs, where schizophrenia samples are mostly hypermethylated compared to control samples (75.9%, 63 out of 83, *P* = 2.4×10^−6^ by binomial test). These trends extend beyond FDR < 0.2 when the top 1,000 DMPs per each cell-type are examined (*P* < 10^−15^ for both cell-types, Binomial tests, Supplementary Fig. 7). In contrast, this bias is not observed in the permuted data (NeuN^+^ empirical *P* = 0.07 and OLIG2^+^ empirical *P* = 0.02, Supplementary Fig. 7).

Considering that NeuN^+^ tend to be more hypermethylated compared to OLIG2^+^ (Fig. 2b), we investigated whether disease patterns in schizophrenia contribute to reduced cell-type difference in DNA methylation. Indeed, szDMPs consistently show decreased cell-type methylation difference compared to control samples (Fig. 4c). In other words, schizophrenia-associated modification of DNA methylation effectively diminishes cell-type distinctive epigenetic profiles in our data.

These results also suggest that sites that did not pass the FDR cutoff but have been detected in the differential methylation analyses may harbor meaningful candidates for future studies. Indeed, genes harboring the top 1,000 szDMPs show enrichment for brain-related functions and diseases (Supplementary Fig. 11). Interestingly, two szDMPs detected in OLIG2^+^ are located within regions reported to be associated with schizophrenia by GWAS^4^ including a CpG located in the intron of *NT5C2* gene, involved in purine metabolism. Indeed, we find that szDMPs tend to co-localize with genetic variants associated with schizophrenia but not with other mental or non-mental traits, mostly with genetic variants below the strict GWAS significance threshold but with moderate-to-high effect sizes (Fig. 4d).

### Cell-type expression differences associated with schizophrenia

Compared to subtle DNA methylation difference, gene expression shows good separation between schizophrenia and control (Fig. 5a) and diagnosis has a strong effect on the variance compared to other covariates (Fig. 5b). We identified 140 and 167 differentially expressed genes between control and schizophrenia (referred to as ‘szDEGs’ henceforth) for NeuN^+^ and OLIG2^+^, respectively at FDR < 0.01 (Fig. 5c; Supplementary Tables 8-9; Methods). We compared our results to previous results obtained from bulk tissues^5, 6^ and identified common and distinct sets of differentially expressed genes across the data sets (Supplementary Tables 10-11; Methods). Comparing effect sizes of commonly differentially expressed genes (*P* < 0.05) among the three data sets analyzed, we find strong correspondence to the CMC and BrainSeq data sets^5, 6^ in NeuN^+^, but not when we compare OLIG2^+^ (Fig. 5d). These results are consistent with the aforementioned deconvolution analysis, indicating that bulk tissue brain studies were limited in terms of non-neuronal signals, such as those coming from oligodendrocytes.

**Figure 5.**
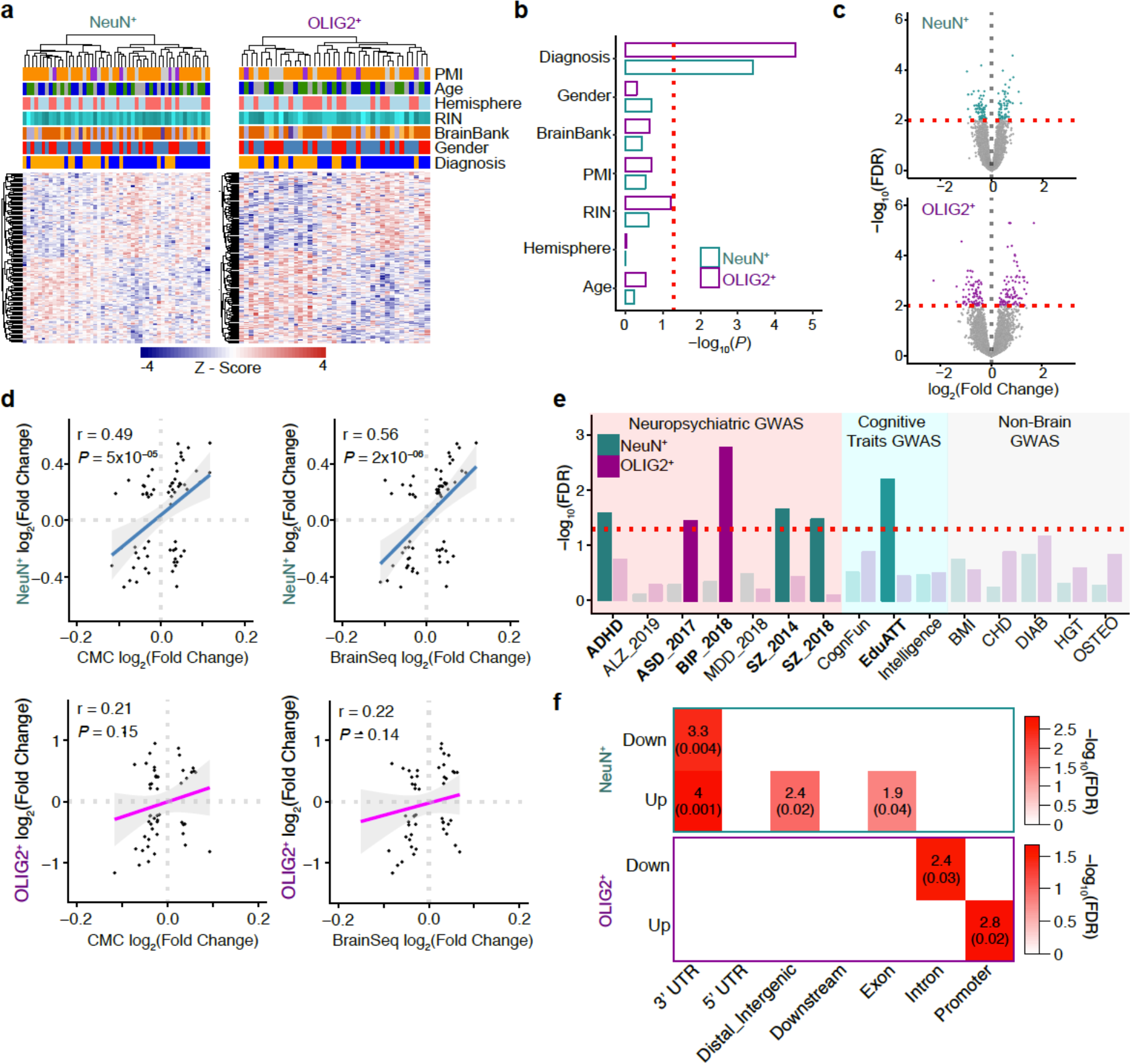
Gene expression associated with schizophrenia in NeuN^+^ and OLIG2^+^. **a.** Heatmap of szDEGs for each cell-type with covariates indicated. **b**. The first principal component of the DEGs was associated with diagnosis. Red dotted line corresponds to *P* = 0.05. **c.** Volcano plot showing szDEGs. X-axis indicates the log_2_(Fold Change), y-axis indicates log_10_(FDR). szDEGs (FDR < 0.01) are colored. **d.** Comparisons of differentially expressed genes in schizophrenia from the current study with the BrainSeq and CMC data. We used genes that are classified as differentially expressed in all three data sets (each dot represents a gene, 63 genes are common to NeuN^+^, CMC and BrainSeq and 49 to OLIG2^+^, CMC, and BrainSeq). The x-axes represent the fold-change between control and schizophrenia in CMC or BrainSeq data sets and the y-axes represent the log_2_(Fold Change) between control and schizophrenia in the current data sets, for either NeuN^+^ specific or OLIG2^+^ specific genes. Regression line and confidence interval are shown for each comparison. **e**. Barplot highlighting the enrichment for trait-associated genetic variants. Bars correspond to NeuN^+^ (cyan) and OLIG2^+^ (magenta) szDEGs. Red dashed line corresponds to the FDR threshold of 0.05. X-axis shows the acronyms for the GWAS data utilized for this analysis (*ADHD:* attention deficit hyperactivity disorder*, ASD:* autism spectrum disorders*, BIP:* bipolar disorder*, ALZ:* Alzheimer’s disease*, MDD:* major depressive disorder, *SZ:* schizophrenia, CognFun: Cognitive Function, EduAtt: Educational Attainment, Intelligence: Intelligence, *BMI:* Body mass index, *CAD:* Coronary Artery Disease, *DIAB:* Diabetes, *HGT:* Height, *OSTEO:* osteoporosis. **f.** Enrichment map for szDEGs (up-/down-regulated) and top 1,000 szDMPs (x-axis shows genic annotation). Enrichment analyses were performed using a Fisher’s Exact Test. Reported odds ratios and FDRs within parentheses for NeuN^+^ (top) and OLIG2^+^ (bottom).

Newly identified szDEGs are enriched for variants for specific disorders or cognitive traits (Fig. 5e; Methods). Notably, NeuN^+^ szDEGs are enriched for GWAS signal from schizophrenia, ADHD as well as educational attainment. Interestingly, OLIG2^+^ szDEGs are enriched for genetic variants associated with bipolar disorder and autism spectrum disorders (Fig. 5e), indicating potential cell-type specific relationship between genetic variants and disease-associated variation of gene expression.

Finally, we investigated the relationship between schizophrenia-associated differential DNA methylation and differential gene expression. Remarkably, similar to what we have observed in DNA methylation, szDEGs are preferentially found in genes that are significantly differentially expressed between cell types for both NeuN^+^ (OR = 7.7, FDR = 8×10^−8^) and OLIG2^+^ (OR = 13, FDR = 7×10^−13^), furthering the functional implication of cell-type specific regulation in schizophrenia. Due to the small number of szDMPs identified at FDR < 0.2, there was little direct overlap between szDMPs and szDEGs. However, when we examined top 1,000 szDMPs, we begin to observe significant enrichments of szDMPs in szDEGs (Fig. 5f). Notably, top 1,000 szDMPs are enriched in genic (3’UTR and exon) and intergenic CpGs in NeuN^+^ (Fig. 5f), while OLIG2^+^ show specific enrichment for intronic and promoter CpGs (Fig. 5f). These results underscore the promise of cell-type specific approaches to elucidate the relationships between genetic variants, epigenetic modifications, and gene expression in a complex neuropsychiatric disorder.

## DISCUSSION

The etiology of schizophrenia remains largely unresolved even though significant efforts have gone into understanding the genetic and molecular mechanisms of the disease^1^. These efforts have been challenged by both the genetic heterogeneity of the disorder as well as the inherent cellular heterogeneity of the brain. To address these issues, we integrated whole-genome sequencing, transcriptome and epigenetic profiles from two major cell-types in the brain. Whole-genome patterns of DNA methylation and gene expression are highly distinct between cell-types, complementing other analyses of cell-type specific epigenetic variation^9, 29^. In particular, our data offer novel resource from oligodendrocytes, a major yet relatively underexplored cell type in the human brains. Indeed we show evidence that previous analyses of bulk tissue gene expression were underpowered to detect oligodendrocyte specific signals, underscoring the strength of a cell-specific approach and the fact that most bulk tissue brain studies tend to focus on or specifically isolate gray matter.

To our knowledge, this is the first study to identify cell-specific correspondence between whole-genome methylation and expression in schizophrenia brains. Compared to substantial cell-type differences, methylation differences between control and schizophrenia are small. Considering 20% false positives, we identified 94 szDMPs, compared to over 4 million cell-type specific DMPs identified at a more stringent cutoff of Bonferroni *P* < 0.05. Nevertheless, schizophrenia-associated epigenetic and transcriptomic alteration is highly cell-type specific, thus offering further support to the idea that cell-type specific regulation may be implicated in schizophrenia pathophysiology^9, 29^. Notably, our resource provides novel whole-genome methylation data from affected brain samples rather than making these connections based on genetic associations. By doing so, we demonstrate that cell-type epigenetic difference is reduced in affected individuals, offering a potential mechanistic link between dysregulation of cell-type specific epigenetic distinction and disease etiology. Similar to the distribution of schizophrenia genetic risk loci, a large portion of differential DNA methylation between control and schizophrenia occur in non-coding regions, which are not accessed by most DNA methylation arrays. We provide detailed interrogation of DNA methylation difference between control and schizophrenia, which can be used for prioritizing targets for further experimental analyses. Even though the nucleotide-resolution information in the current study is counterbalanced by the relatively limited number of samples (*N*=45 and 50 from control and schizophrenia brains), with rapidly decreasing sequencing costs, future WGBS studies with increased sample sizes are sure to follow and will allow us to elucidate biological alteration associated with schizophrenia.

## Acknowledgements

We thank the donors and their families for the tissue samples used in these studies. We also thank Angela Mobley of the Flow Cytometry Facility and Vanessa Schmid of the Next Generation Sequencing Core of UT Southwestern Medical Center for technical support. G. K. is a Jon Heighten Scholar in Autism Research at UT Southwestern. This work was supported by the Uehara Memorial Foundation to N.U.; JSPS Grant-in-Aid for Early-Career Scientists (18K14814) to N.U. and Scientific Research (C) (18K06977) to K.T.; Takeda Science Foundation to N.U.; the JSPS Program for Advancing Strategic International Networks to Accelerate the Circulation of Talented Researchers (S2603) to S.B., N.U., K.T. and G.K.; the James S. McDonnell Foundation 21^st^ Century Science Initiative in Understanding Human Cognition – Scholar Award to G.K.; National Science Foundation (SBE-131719) to S.V.Y; the National Chimpanzee Brain Resource, NIH R24NS092988, the NIH National Center for Research Resources P51RR165 (superceded by the Office of Research Infrastructure Programs/OD P51OD11132) to T.M.P; and the NIMH (MH103517), to T.M.P., G.K., and S.V.Y. Human tissue samples were obtained from the NIH NeuroBioBank (The Harvard Brain Tissue Resource Center, funded through HHSN-271-2013-00030C; the Human Brain and Spinal Fluid Resource Center, VA West Los Angeles Healthcare Center; and the University of Miami Brain Endowment Bank) and the UT Psychiatry Psychosis Research Program (Dallas Brain Collection).

## Author Contributions

I.M., S.B., G.K., and S.V.Y. analyzed the data and wrote the paper. N.U. and K.T. designed, optimized and performed the nuclei Isolations, nuclei sorting, gDNA and RNA preparations, and RNA-seq. C.D. performed additional cell sorting and gDNA extraction. P.C. performed WGS and WGBS. T.L. performed validation experiments. I.M. analyzed WGBS and WGS data. S.B. analyzed RNA-seq data. I.H. constructed linear mixed effect models and provided statistical guidance. H.J. analyzed validation and hydroxymethylation data sets. C.T. provided human post-mortem brain tissue. T.M.P., G.K., and S.V.Y. designed and supervised the study, and provided intellectual guidance. All authors discussed the results and commented on the manuscript.

## Online Methods

### Sampling strategy

Frozen brain specimens from Brodmann area 46 were obtained from several brain banks (Supplementary Tables 1-2). Cases and controls were matched by age group and additional demographics such as gender were matched when possible (Supplementary Table 1). Information on comorbidities and cause of death when known are included in Supplementary Table 1.

### Nuclei Isolation from human post-mortem brain

Nuclei Isolation was performed as described previously^18, 34^ with some modifications. Approximately 700 mg of frozen postmortem brain was homogenized with lysis buffer (0.32 M sucrose, 5 mM CaCl_2_, 3 mM Mg(Ac)_2_, 0.1 mM EDTA, 10 mM Tris-HCl pH8.0, 0.1 mM PMSF, 0.1% (w/o) Triton X-100, 0.1% (w/o) NP-40, protease inhibitors (1:100) (#P8340, Sigma, St. Louis, MO), RNase inhibitors (1:200) (#AM2696, ThermoFisher, Waltham, MA)) using a dounce homogenizer. Brain lysate was placed on a sucrose solution (1.8 M sucrose, 3 mM Mg(Ac)_2_, 10 mM Tris-HCl pH8.0) to create a concentration gradient. After ultracentrifuge at 24,400 rpm for 2.5 hours at 4°C, the upper layer of the supernatant was collected as the cytoplasmic fraction. The pellet, which included nuclei, was resuspended with ice-cold PBS containing RNase inhibitors, and incubated with mouse alexa488 conjugated anti-NeuN (1:200) (#MAB377X, Millipore, Billerica, MA) and rabbit alexa555 conjugated anti-OLIG2 (1:75) (#AB9610-AF555, Millipore) antibodies with 0.5% BSA for 45 min at 4°C. Immuno-labeled nuclei were collected as NeuN-positive or OLIG2-positive populations by fluorescence-activated nuclei sorting (FANS). After sorting, gDNA and total RNA were purified from each nuclei population using a ZR-Duet DNA/RNA MiniPrep (Plus) kit (#D7003, Zymo Research, Irvine, CA) according to the manufacturer’s instruction. Total RNA was treated with DNase I after separation from gDNA. 200 ng total RNA from each sample was treated for ribosomal RNA removal using the Low Input RiboMinus Eukaryote System v2 (#A15027, ThermoFisher) according to the manufacturer’s instruction. After these purification steps, gDNA and total RNA were quantified by Qubit dsDNA HS (#Q32851, ThermoFisher) and RNA HS assay (#Q32852, ThermoFisher) kits, respectively. Immunostaining was visualized using a Zeiss LSM 880 with Airyscan Confocal Laser Scanning Microscope. 100 µl of sorted nuclei were placed onto microscope slides, 300 µl of ProLong Diamond Antifade Mountant with DAPI (#P36971, ThermoFisher) was added, and covered with glass coverslips before imaging.

### Whole-genome bisulfite sequencing (WGBS) library generation and data processing

As a control for bisulfite conversion, 10 ng of unmethylated lambda phage DNA (#D1501, Promega) was added to the 1 µg of input DNA. Libraries were made with an in-house Illumina sequencer compatible protocol. The extracted DNA was fragmented by S-series Focused-ultrasonicator (Covaris, Woburn, MA) using the “200 bp-target peak size protocol”. Fragmented DNA was then size selected (200-600 bp) with an Agencourt AMPure XP bead-based (#A63880, Beckman Coulter, Brea, CA) size selection protocol^35^. The DNA End repair step was performed with End-It DNA End-Repair Kit (#ER81050, Epicentre, Madison, WI). After End repair step, A-tailing (#M0202, New England Biolabs, Ipswich, MA) and ligation steps were performed to ligate methylated adaptors.

Bisulfite treatment of gDNA was performed using the MethylCode Bisulfite Conversion Kit (#MECOV50, ThermoFisher). Purified gDNA was treated with CT conversion reagent in a thermocycler for 10 min at 98°C, followed by 2.5 hours at 640°C. Bisulfite-treated DNA fragments remain single-stranded as they are no longer complementary. Low-cycle (4-8) PCR amplification was performed with Kapa HiFi Uracil Hotstart polymerase enzyme (#KK2801, KAPA Biosystems, Wilmington, MA) which can tolerate Uracil residues. The final library fragments contain thymines and cytosines in place of the original unmethylated cytosine and methylated cytosines, respectively.

The methylome libraries were diluted and loaded onto an Illumina HiSeq 2500 or HiSeqX system for sequencing using 150 bp paired-end reads. We generated over 900 million reads per sample and performed quality and adapter trimming using TrimGalore v.0.4.1 (Babraham Institute) with default parameters. Reads were mapped first to the PhiX genome to remove the spike-in control and the remaining reads were mapped to the human GRCh37 (build 37.3) reference genome using Bismark v 0.14.5^36^ and bowtie v1.1.2^37^. We removed reads with exact start and end positions using Bismkar deduplication script. After de-duplication, we calculated fractional methylation levels at individual cytosines^27^. Overall, we generated a total of 72.6 billion reads (equivalent to 10.9 tera base pairs of raw sequence data) and obtained per-sample average coverage depths >25× covering 98% of the 28 million CpGs in the human genome (Supplementary Table 12). Bisulfite conversion rates were estimated by mapping the reads to the lambda phage genome (NC_001416.1). See Supplementary Fig. 12 for a general overview of WGBS data quality and processing.

### Whole-genome sequencing (WGS) data processing

Quality and adapter trimming were performed using TrimGalore v.0.4.1 (Babraham Institute) with default parameters. Reads were mapped to the human GRCh37 reference genome using BWA v0.7.4^38^ and duplicates were removed using picard v2.8.3 (https://broadinstitute.github.io/picard/index.html). We identified genetic polymorphisms from re-sequencing data following GATK v3.7 best practices workflow^39^. Specifically, we used HapMap 3.3, Omni 2.5M, 1,000 Genomes Phase I and dbSNP 138 as training data sets for variant recalibration. We filtered variant calls with high genotype quality (GQ >=20.0). Overall, we generated a total of 225 million reads and identified 15,331,100 SNPs with mean depth above >16.5× (Supplementary Table 13). We removed polymorphic cytosines from downstream differential methylation analyses keeping a total of 24,942,405 autosomal CpGs (Supplementary Table 14). See Supplementary Fig. 12 for a general overview of WGS data quality and processing.

For quality control of the SNP calling, we performed principal component analyses using an additional 210 samples from 4 different populations from the HapMap Project (60 CEU, 90 CBH/JPT and 60 YRI) to explore the genetic ancestry of the individuals. After LD pruning (r^2^>0.2) with SNPRelate R package, we used 66,667 autosomal polymorphic SNPs in the analysis. The PC plot shows that the reported ancestry of the individuals was mostly concordant to that inferred from the SNPs (Supplementary Fig. 13), validating the genotype calling. The first 10 genetic PCs were included in the differential methylation analyses to control for population structure (Supplementary Table 14).

### Hierarchical clustering of methylomes from diverse human cell-types

We added WGBS data from additional tissues^12^ (see original references for the data sets therein) and Lister et al.^23^, and the corresponding genome coordinates (hg38 and hg18) were converted to hg19 using UCSC Batch Coordinate Conversion tool (liftOver executable)^40^. The sample indicated with the star in Figure 2a was also remapped to hg38 from raw data following the same protocol as other non-brain tissues (from Mendizabal and Yi, 2016) and lifted over to hg19. The clustering of the two methylomes from the same individual “NeuN+_ind2” suggests no significant effect of mapping/lift over in the clustering results. A total of 14,115,607 CpG positions with at least 5× coverage in all individuals were used to draw a hierarchical clustering tree (using R stats package’s hclust function with method=average (=UPGMA) based on Euclidean distances using fractional methylation values using dist function). The tree was plotted using dendextend and circlize packages.

### Identification of differentially methylated positions (DMPs) and regions (DMRs) between OLIG2^+^ and NeuN^+^

We identified DMPs between 25 NeuN^+^ and 20 OLIG2^+^ individuals by using DSS^24^. DSS handles variance across biological replicates as well as models read counts from WGBS experiments. Importantly, DSS also considers other biological covariates that may affect DNA methylation patterns. Specifically, we considered age, gender, brain hemisphere, post-mortem interval (PMI), conversion rates, brain bank, and genetic ancestry (using the first 10 genetic PCs obtained from WGS of the same individuals) as covariates (Supplementary Tables 1-2 and 14, and Supplementary Fig. 14). Age and PMI were converted to categorical variables (“AgeClass” and “PMIClass” in Supplementary Table 2).

Since C>T and G>A polymorphisms at CpGs could generate spurious differentially methylated sites on bisulfite conversion experiments, we excluded polymorphic CpGs (identified from re-sequencing the same panel of individuals, Supplementary Table 15) from DMP analyses. For DMP identification between OLIG2^+^ and NeuN^+^ samples, we used a Bonferroni cutoff on *P* < 0.05 and identified 4,058,898 DMPs out of 24,596,850 CpGs tested. For DMR identification, we considered a minimum region of 50 bp with at least 5 significant DMPs and identified 145,073 regions (Supplementary Table 3). We explored the effect of coverage on cell-type DMP identification and found that low coverage sites had a limited contribution to the significant DMPs, indeed relatively more sites were detected at more stringent coverage thresholds. For example, removing sites <5× in 80% of individuals within each cell-type, led to a total of 4,037,979 significant DMPs at Bonferroni 0.05 cutoff (out of 23,788,847 CpGs, 16.97%), whereas the removal of sites <10× lead to 3,903,652 DMPs (out of 21,399,153 CpGs tested, 18.2%), and <20× lead to 2,509,489 DMPs (out of 10,960,268 CpGs considered, 23.8%). Enrichment between cell-type DMPs and szDMP, and between cell-type DMPs and ChIP-seq peaks were similar when using the >20× coverage data sets instead of using all sites.

Of note, as our differential methylation analyses are run under a multifactor design in DSS, the estimated coefficients in the regression are based on a generalized linear model framework using arcsine link function to reduce dependence of variance on the fractional methylation levels^24, 41^. Thus, whereas the direction of change is indicated by the sign of the test statistic, its values cannot be interpreted directly as fractional methylation level differences. The distribution of the statistic depends on the differences in methylation levels and biological variations, as well as technical factors such as coverage depth. For DMRs, the method provides “areaStat” values which is defined as the sum of the test statistic of all CpG sites within the DMR. To obtain a more interpretable estimate of fractional methylation differences we also provide results for a linear model using the same formula as for DSS.

### Functional characterization of DMRs

For different enrichment analyses, we generated matched control regions. We generated 100 sets of regions with similar genomic properties as the DMRs: number of total regions, region length distribution, chromosome, and matched GC content within 1%. Empirical *P* values were computed by counting the number of matched control sets showing values as extreme as the observed one. Enrichments were computed as the ratio between the observed value and the mean of the matched control sets. We used ChIPSeeker^42^ and bioconductor’s UCSC gene annotation library TxDb.Hsapiens.UCSC.hg19.knownGene to annotate DMRs to genes. We explored the 25 chromatin state-model maps based on ChIP-Seq experiments on 6 chromatin marks (H3K4me3, H3K4me1, H3K36me3, H3K27me3, H3K9me3 and H3K27ac) from the Roadmap Epigenomics Project^28^. We joined several categories related to enhancer states, including TxReg, TxEnh5’, TxEnh3’, TxEnhW, EnhA1, EnhA2, EnhW1, EnhW2, EnhAc.

### Overlap with neuronal and non-neuronal ChIP-seq data sets

We analyzed the overlap between our cell-type specific DMPs and DMRs with neuron and non-neuron histone mark data on H3K4me3 and H3k27ac ChIP-seq experiments^9^. We only considered peaks that were assigned as “neuronal” and “non-neuronal” and discarded “NS” peaks from the Supplementary Table 11 in the cited paper. To test directionality with our OLIG2^+^ vs. NeuN^+^ differentially methylated sites, we further discarded peaks that overlapped between cell-types (i.e. neuronal H3K4me3 peaks overlapping with non-neuronal H3K27ac, and non-neuronal H3K4me3 peaks overlapping with neuronal H3K27ac peaks).

### Non-CpG methylation patterns in brain cell-types

We studied DNA methylation patterns of NeuN^+^ and OLIG2^+^ outside CpG dinucleotides (CH context). Given the low fractional patterns of DNA methylation outside CpG sites, and to minimize the influence of any additional covariates, only individuals with conversion rates >=0.995 were considered (15 NeuN^+^ and 14 OLIG2^+^). We filtered cytosines that showed less than 5× coverage in 90% of individuals per cell-type, as well as removed positions with genetic polymorphisms (C>T and T>C SNPs to account for SNPs at both strands). A total of 333 and 457 million cytosines remained in NeuN^+^ and OLIG2^+^, respectively. Cytosines in gene bodies were filtered using BEDtools^43^.

### Identification of DMPs between schizophrenia and control individuals

We used DSS to identify DMPs between schizophrenia and control samples. Again, we considered biological covariates in the differential methylation analyses, namely age, gender, brain hemisphere, PMI, conversion rates, brain bank, and genetic ancestry (using the first 10 genetic PCs obtained from WGS of the same individuals, see File S3 for specific commands used). Using all sites without minimum coverage filtering (24,596,850 CpGS) and an FDR cutoff of 0.2 for significance identified a total of 201 and 60 DMPs in OLIG2^+^ and NeuN^+^ respectively. Given the overall modest differences between cases and controls, we further filtered sites with less than 20× in at < 80% of individuals per group. A total of 7,682,148 CpGs and 11,963,682 CpGs remained in NeuN^+^ and OLIG2^+^ data sets respectively. By applying an FDR cutoff of 0.2 for significance, we identified 14 and 83 significant DMPs in NeuN^+^ and OLIG2^+^, respectively.

As a comparison, we also ran differential methylation analyses for disease using a linear model based on fractional methylation values for every CpGs site, and considered the same covariates as in the DSS analyses. We plotted quantile-quantile plots for the expected and observed *P* values obtained from DSS and linear model analyses between schizophrenia and control, as well as to evaluate how coverage affects these two methods. We observed that DSS provides correction for low coverage sites. Note the systematic depletion of good *P* values at low coverage sites in DSS (Supplementary Fig. 15), compared to high coverage sites. In contrast, a linear model shows similar genome-wide distribution of *P* values at low and high coverage sites. We identified a total of 60 and 210 CpGs in NeuN^+^ and OLIG2^+^ respectively at FDR < 0.2. However, to obtain a more conservative set of hits, we additionally filter for high coverage sites (20× in at least 80% of samples per disease-control group) and recalculated FDR, obtaining 14 and 83 significant sites at FDR < 0.2. In order to test the robustness of the results and the effect of covariates as well as potential hidden structures in the data, we performed a permuting analysis by randomly assigning case/control labels and re-ran DSS 100 times.

### szDMP gene annotation and functional enrichment

We used ChIPSeeker^42^ and bioconductor’s UCSC gene annotation library TxDb.Hsapiens.UCSC.hg19.knownGene to annotate top 1,000 szDMPs to genes (ordered by *P* values). We used genes associated to genic szDMPs only (all annotation categories excluding distal intergenic, defined as >1.5kb from start or end of genes) for functional enrichment using ToppGene^44^.

### szDMP enrichment at GWAS

Genome-wide *P* values and odds-ratios for GWAS for schizophrenia^4^, smoking^45^, clozapine induced agranulocytosis^46^, coronary artery disease, bipolar disorder^47^, autism spectrum disorder, anorexia nervosa, were downloaded from the Psychiatric Genomics Consortium at https://www.med.unc.edu/pgc/files/resultfiles/. Data for rheumatoid arthritis^48^ were downloaded from ftp://ftp.broadinstitute.org/pub/rheumatoid_arthritis/Stahl_etal_2010NG/. In order to quantify the differences between traits (in particular at SNPs below the significance thresholds, where the majority of the schizophrenia heritability is found), we followed the following approach. For each szDMP, we identified all SNPs reported by each GWAS study within a 1kb window and kept the number of SNPs at different quantiles of absolute odds ratio (OR). We repeated this step using the same number of random non-DMPs 100 times. To obtain empirical *P* values, we calculated the number of times non-szDMP sets showed more SNPs in each OR quantile than szDMPs. SNPs showing moderate-to-high OR in schizophrenia GWAS consistently showed low empirical *P* values for both cell-type DMPs, implying that SNPs close to szDMPs show bigger effect sizes.

### Hydroxymethylation at szDMPs

We compared our results to a single-base resolution hydroxymethylome maps^49^. Specifically, TAB-seq data from an adult human brain sample was obtained from GEO (GSE46710). We used the sites presenting high hmC as defined in the original paper (hmC > mC; *n* = 5,692,354). We plotted quantile-quantile plots of DSS statistic *P* values at high hmC loci and random loci. These analyses showed no significant presence of hmC in the szDMPs (Supplementary Fig. 16).

### Smoking DMPs at szDMP

We explored the co-localization of szDMPs with CpGs associated with tobacco smoking^50–52^. None of the analyzed smoking DMPs (n=206) were found among our szDMPs at FDR < 0.2 nor at top 1,000 CpGs with best *P* values per cell-type. These analyses suggest that szDMPs might not be confounded by smoking.

### Targeted validation experiments

We designed high coverage bisulfite experiments to sequence 18 regions (Supplementary Table 16) from 44 samples (including 24 new individuals not included in the WGBS experiments, Supplementary Table 17). We conducted bisulfite conversions of gDNA from OLIG2^+^ and NeuN^+^ cells using EZ DNA Methylation-Gold Kit (#D5006, Zymo Research) according to the manufacturer’s instructions. Sodium bisulfite converted unmethylated cytosines to uracil while methylated cytosines remained unconverted. Upon subsequent PCR amplification, uracil was ultimately converted to thymine. Bisulfite-sequencing PCR primers were designed using MethPrimer 2.0 and BiSearch to target a panel of 12 loci in OLIG2^+^ and 6 loci in NeuN^+^ (Supplementary Table 16). The primers were designed with an Illumina adaptor overhang. The sites of interest were amplified using JumpStart Taq DNA polymerase (#D9307, Sigma) and quantified using gel electrophoresis to verify size and Qubit fluorometric quantitation to determine concentration. Equimolar quantities of each of the target amplicons were pooled for each individual and NGS libraries were prepared in a second PCR reaction according to Nextera XT DNA Sample Preparation protocol. Libraries were barcoded with a unique pair of Nextera XT primers. The libraries were sequenced with Illumina MiSeq using the 500-cycle kit (250 paired end sequencing). We sequenced the samples at high coverage using a MiSeq machine and 250bp paired end reads at the Georgia Institute of Technology High Throughput DNA Sequencing Core. We mapped the reads to the human GRCh37 (build 37.3) reference genome using Bismark v0.20.2 and Bowtie v2.3.4. We trimmed the reads for low quality and adapters using TrimGalore v.0.5.0 (Babraham Institute) with default parameters. Only sites with at least 200× coverage were considered (mean=14,580, median=10,810). One region showed low read counts and was excluded (Supplementary Table 16). A total of 16 DMPs and an additional 50 adjacent CpGs were considered in the validation analyses. Fractional methylation values were adjusted for covariates using the following linear model: lm (methylation ~ Diagnosis + Sex + Age_Class + PMI_Class).

### Concordance with previous methylation studies on schizophrenia

We evaluated the concordance between our disease DMP signals with Jaffe et al.^7^ which used bulk brain tissue and Illumina 450K chips. We binned Jaffe et al study’s whole-genome *P* values and calculated the fraction of CpGs in our study showing the same directionality in both studies (i.e. hypomethylated or hypermethylated in disease vs control). For each cell-type, we tested the significance at each *P* value bin using a Binomial test with *P* = 0.5 expectation. We additionally compared the distribution of concordance rates from the 100 control data sets obtained using case/control permuted labels and re-running DSS on them.

### RNA-sequencing (RNA-seq)

RNA-seq was performed as described previously^53^. Total RNA from the cytoplasmic fraction was extracted with the miRNeasy Mini kit (#217004, Qiagen, Hilden, Germany) according to the manufacturer’s instruction. The RNA integrity number (RIN) of total RNA was quantified by Agilent 2100 Bioanalyzer using Agilent RNA 6000 Nano Kit (#5067-1511, Agilent, Santa Clara, CA). Total RNAs with an average RIN value of 7.5±0.16 were used for RNA-seq library preparation. 50 ng of total RNA after rRNA removal was subjected to fragmentation, first and second strand syntheses, and clean up by EpiNext beads (#P1063, EpiGentek, Farmingdale, NY). Second strand cDNA was adenylated, ligated and cleaned up twice by EpiNext beads. cDNA libraries were amplified by PCR, and cleaned up twice by EpiNext beads. cDNA library quality was quantified by a 2100 Bioanalyzer using an Agilent High Sensitivity DNA Kit (#5067-4626, Agilent). Barcoded libraries were pooled and underwent 75 bp single-end sequencing on an Illumina NextSeq 500.

### RNA-seq mapping, QC and expression quantification

Reads were aligned to the human hg19 (GRCh37) reference genome using STAR 2.5.2b^54^ with the following parameters: “*--outFilterMultimapNmax 10 --alignSJoverhangMin 10 -- alignSJDBoverhangMin 1 --outFilterMismatchNmax 3 --twopassMode Basic*”. Ensemble annotation for hg19 (version GRCh37.87) was used as reference to build STAR indexes and alignment annotation. For each sample, a BAM file including mapped and unmapped reads with spanning splice junctions was produced. Secondary alignment and multi-mapped reads were further removed using in-house scripts. Only uniquely mapped reads were retained for further analyses. Quality control metrics were performed using RseqQC using the hg19 gene model provided^55^. These steps include: number of reads after multiple-step filtering, ribosomal RNA reads depletion, and defining reads mapped to exons, UTRs, and intronic regions. Picard tool was implemented to refine the QC metrics (http://broadinstitute.github.io/picard/). Gene level expression was calculated using HTseq version 0.9.1 using intersection-strict mode by Exons^56^. Counts were calculated based on protein-coding genes annotation from the Ensemble GRCh37.87 annotation file. See quality control metrics in Supplementary Fig. 17-18 and Supplementary Table 18.

### Covariate adjustment and differential expression

Counts were normalized using counts per million reads (CPM). Genes with no reads in either schizophrenia (SZ) or control (CTL) samples were removed. Normalized data were assessed for effects from known biological covariates (diagnosis, age, gender, hemisphere), technical variables related to sample processing (RIN, brain bank, PMI), and technical variables related to surrogate variation (SV) (Supplementary Fig. 19). SVs were calculated using SVA^57^ based on “be” method with 100 iterations. The data were adjusted for technical covariates using a linear model:

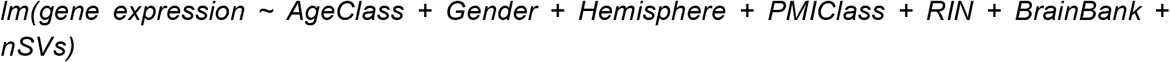

Adjusted CPM values were used for co-expression analysis and visualization. For differential expression, we used the lmTest ("robust") and ebayes functions in the limma^58^ fitting all of the statistical models to estimate log_2_ fold changes, *P* values and FDR/Bonferroni correction. This method was use for 1) cell-type differences (|log_2_(Fold Change)| > 0.5 and Bonferroni FDR < 0.05), 2) NeuN^+^ SZ-CTL analysis (|log_2_(Fold Change)| > 0.3 and FDR < 0.01), 3) OLIG2^+^ SZ-CTL analysis (|log_2_(Fold Change)| > 0.3 and FDR < 0.01). Bonferroni was used in 1 to provide higher stringency on the data analysis.

### Cross-validation

Cross-validation analyses were applied to ensure robustness of the DEG analysis:

1. Permutation method based on gene expression randomization (nPerm=200).
2. Leave-one-out method based on subsampling the data (nLOO = 200).

### Functional gene annotation

The functional annotation of differentially expressed and co-expressed genes was performed using ToppGene^44^. A Benjamini-Hochberg FDR (*P* < 0.05) was applied as a multiple comparisons adjustment.

### GWAS data and enrichment

We manually compiled a set of GWAS studies for several neuropsychiatric disorders, cognitive traits and non-brain disorders/traits. Summary statistics from the genetic data were downloaded from Psychiatric Genomics Consortium (http://www.med.unc.edu/pgc/results-and-downloads) and GIANT consortium (https://portals.broadinstitute.org/collaboration/giant/). Gene level analysis was performed using MAGMA^59^ v1.04, which considers linkage disequilibrium between SNPs. 1,000 Genomes (EU) data set was used as reference for linkage disequilibrium. SNPs annotation was based on the hg19 genome annotation (gencode.v19.annotation.gtf). MAGMA statistics and –log10(FDR) are reported in Supplementary Table 19 for each of the GWAS data analyzed. Brain GWAS: *ADHD:* attention deficit hyperactivity disorder^60^*, ASD:* autism spectrum disorders (https://www.biorxiv.org/content/early/2017/11/27/224774), BIP: bipolar disorder^61^*, ALZ:* Alzheimer’s disease^62^*, MDD:* major depressive disorder^63^, *SZ:* schizophrenia^4, 61^. Cognitive Traits GWAS: CognFun = Cognitive Function ^60^, EduAtt = Educational Attainment^64^, Intelligence = Intelligence^65^. Non-Brain GWAS: *BMI* = Body mass index^66^, *CAD* = Coronary Artery Disease^67^, *DIAB* = Diabetes^68^, *HGT* = Height (https://www.biorxiv.org/content/early/2018/07/09/355057), *OSTEO* = osteoporosis^69^.

### Cell-type enrichment and deconvolution analyses

MTG single-nuclei RNA-seq was downloaded from Allen Brain institute web-portal^70^). Normalized data and cluster annotation were used to define cell-markers using FindAllMarkers in Seurat^71^ with the following parameters: logfc.threshold = 0.25, test.use = "wilcox", min.pct = 0.25, only.pos = TRUE, return.thresh = 0.01, min.cells.gene = 3, min.cells.group = 3. Enrichment analyses were performed using a Fisher’s Exact Test. Cell-type deconvolution was performed using MuSiC^72^ with the following parameters: iter.max = 1000, nu = 1e-10, eps = 0.01, normalize=F.

### Public data analyses

GTEx tissue expression was downloaded from the GTEx web-portal. Raw data was normalized using log_2_(CPM+1)^73^. Gene expression data from SZ and healthy CTL brain tissue was downloaded from the Common Mind Consortium^5^. Gene expression data from SZ and healthy CTL developmental brain tissue was downloaded from Brain Phase1^6^. We applied differential expression analysis using the lmTest ("robust") and ebayes functions in the limma^58^ fitting all of the technical/biological covariates and surrogate variables to estimate log2 fold changes, *P* values and FDR/Bonferroni correction. Surrogate variables were calculated with SVA package^57^.

## References

1. Birnbaum, R. & Weinberger, D.R. Genetic insights into the neurodevelopmental origins of schizophrenia. Nature Reviews Neuroscience 18, 727 (2017).

2. Gusev, A., et al. Transcriptome-wide association study of schizophrenia and chromatin activity yields mechanistic disease insights. Nat Genet 50, 538–548 (2018).

3. Loh, P.R., et al. Contrasting genetic architectures of schizophrenia and other complex diseases using fast variance-components analysis. Nat Genet 47, 1385–1392 (2015).

4. Schizophrenia Working Group of the Psychiatric Genomics, C. Biological insights from 108 schizophrenia-associated genetic loci. Nature 511, 421–427 (2014).

5. Fromer, M., et al. Gene expression elucidates functional impact of polygenic risk for schizophrenia. Nat Neurosci 19, 1442–1453 (2016).

6. Jaffe, A.E., et al. Developmental and genetic regulation of the human cortex transcriptome illuminate schizophrenia pathogenesis. Nature Neuroscience 21, 1117–1125 (2018).

7. Jaffe, A.E., et al. Mapping DNA methylation across development, genotype and schizophrenia in the human frontal cortex. Nat Neurosci 19, 40–47 (2016).

8. Jakovcevski, M. & Akbarian, S. Epigenetic mechanisms in neurological disease. Nature Medicine 18, 1194 (2012).

9. Girdhar, K., et al. Cell-specific histone modification maps in the human frontal lobe link schizophrenia risk to the neuronal epigenome. Nat Neurosci 21, 1126–1136 (2018).

10. Hannon, E., et al. Methylation QTLs in the developing brain and their enrichment in schizophrenia risk loci. Nature Neuroscience 19, 48–54 (2016).

11. Ziller, M.J., et al. Charting a dynamic DNA methylation landscape of the human genome. Nature 500, 477 (2013).

12. Mendizabal, I. & Yi, S.V. Whole-genome bisulfite sequencing maps from multiple human tissues reveal novel CpG islands associated with tissue-specific regulation. Hum Mol Genet 25, 69–82 (2016).

13. Darmanis, S., et al. A survey of human brain transcriptome diversity at the single cell level. Proceedings of the National Academy of Sciences 112, 7285 (2015).

14. Lake, B.B., et al. Neuronal subtypes and diversity revealed by single-nucleus RNA sequencing of the human brain. Science 352, 1586–1590 (2016).

15. Pollen, Alex A., et al. Molecular Identity of Human Outer Radial Glia during Cortical Development. Cell 163, 55–67 (2015).

16. Ecker, J.R., et al. The BRAIN Initiative Cell Census Consortium: Lessons Learned toward Generating a Comprehensive Brain Cell Atlas. Neuron 96, 542–557 (2017).

17. Kozlenkov, A., et al. Differences in DNA methylation between human neuronal and glial cells are concentrated in enhancers and non-CpG sites. Nucleic Acids Research (2013).

18. Jiang, Y., Matevossian, A., Huang, H.S., Straubhaar, J. & Akbarian, S. Isolation of neuronal chromatin from brain tissue. BMC Neurosci 9, 42 (2008).

19. Mighdoll, M.I., Tao, R., Kleinman, J.E. & Hyde, T.M. Myelin, myelin-related disorders, and psychosis. Schizophr Res 161, 85–93 (2015).

20. Hoffman, G.E., et al. Transcriptional signatures of schizophrenia in hiPSC-derived NPCs and neurons are concordant with post-mortem adult brains. Nat Commun 8, 2225 (2017).

21. Micu, I., Plemel, J.R., Caprariello, A.V., Nave, K.A. & Stys, P.K. Axo-myelinic neurotransmission: a novel mode of cell signalling in the central nervous system. Nat Rev Neurosci 19, 49–58 (2018).

22. Birnbaum, R., et al. Investigation of the Prenatal Expression Patterns of 108 Schizophrenia-Associated Genetic Loci. Biological Psychiatry 77, e43–e51 (2015).

23. Lister, R., et al. Global epigenomic reconfiguration during mammalian brain development. Science 341, 1237905 (2013).

24. Park, Y. & Wu, H. Differential methylation analysis for BS-seq data under general experimental design. Bioinformatics 32, 1446–1453 (2016).

25. Hansen, K.D., Langmead, B. & Irizarry, R.A. BSmooth: from whole genome bisulfite sequencing reads to differentially methylated regions. Genome Biol 13, R83 (2012).

26. Huh, I., Yang, X., Park, T. & Yi, S.V. Bis-class: a new classification tool of methylation status using Bayes classifier and local methylation information. BMC Genomics 15, 608 (2014).

27. Lister, R., et al. Human DNA methylomes at base resolution show widespread epigenomic differences. Nature 462, 315–322 (2009).

28. Roadmap Epigenomics, C., et al. Integrative analysis of 111 reference human epigenomes. Nature 518, 317–330 (2015).

29. Rizzardi, L.F., et al. Neuronal brain-region-specific DNA methylation and chromatin accessibility are associated with neuropsychiatric trait heritability. Nature Neuroscience 22, 307–316 (2019).

30. Sarnat, H.B., Nochlin, D. & Born, D.E. Neuronal nuclear antigen (NeuN): a marker of neuronal maturation in early human fetal nervous system. Brain & development 20, 88–94 (1998).

31. Gusel’nikova, V.V. & Korzhevskiy, D.E. NeuN As a Neuronal Nuclear Antigen and Neuron Differentiation Marker. Acta naturae 7, 42–47 (2015).

32. Yokoo, H., et al. Anti-human Olig2 antibody as a useful immunohistochemical marker of normal oligodendrocytes and gliomas. The American journal of pathology 164, 1717–1725 (2004).

33. Rhee, W., et al. Quantitative analysis of mitotic Olig2 cells in adult human brain and gliomas: implications for glioma histogenesis and biology. Glia 57, 510–523 (2009).

34. Matevossian, A. & Akbarian, S. Neuronal nuclei isolation from human postmortem brain tissue. J Vis Exp (2008).

35. Urich, M.A., Nery, J.R., Lister, R., Schmitz, R.J. & Ecker, J.R. MethylC-seq library preparation for base-resolution whole-genome bisulfite sequencing. Nat Protoc 10, 475–483 (2015).

36. Krueger, F. & Andrews, S.R. Bismark: a flexible aligner and methylation caller for Bisulfite-Seq applications. Bioinformatics 27, 1571–1572 (2011).

37. Langmead, B., Trapnell, C., Pop, M. & Salzberg, S.L. Ultrafast and memory-efficient alignment of short DNA sequences to the human genome. Genome Biol 10, R25 (2009).

38. Li, H. & Durbin, R. Fast and accurate short read alignment with Burrows-Wheeler transform. Bioinformatics 25, 1754–1760 (2009).

39. McKenna, A., et al. The Genome Analysis Toolkit: a MapReduce framework for analyzing next-generation DNA sequencing data. Genome Res 20, 1297–1303 (2010).

40. Kent, W.J., et al. The human genome browser at UCSC. Genome Res 12, 996–1006 (2002).

41. Guan, Y. Variance stabilizing transformations of Poisson, binomial and negative binomial distributions. Stat Probabil Lett 79, 1621–1629 (2009).

42. Yu, G., Wang, L.-G. & He, Q.-Y. ChIPseeker: an R/Bioconductor package for ChIP peak annotation, comparison and visualization. Bioinformatics 31, 2382–2383 (2015).

43. Quinlan, A.R. & Hall, I.M. BEDTools: a flexible suite of utilities for comparing genomic features. Bioinformatics 26, 841–842 (2010).

44. Chen, J., Bardes, E.E., Aronow, B.J. & Jegga, A.G. ToppGene Suite for gene list enrichment analysis and candidate gene prioritization. Nucleic Acids Res 37, W305–311 (2009).

45. Tobacco & Genetics C. Genome-wide meta-analyses identify multiple loci associated with smoking behavior. Nat Genet 42, 441–447 (2010).

46. Goldstein, J.I., et al. Clozapine-induced agranulocytosis is associated with rare HLA-DQB1 and HLA-B alleles. Nat Commun 5, 4757 (2014).

47. Psychiatric, G.C.B.D.W.G. Large-scale genome-wide association analysis of bipolar disorder identifies a new susceptibility locus near ODZ4. Nat Genet 43, 977–983 (2011).

48. Stahl, E.A., et al. Genome-wide association study meta-analysis identifies seven new rheumatoid arthritis risk loci. Nat Genet 42, 508–514 (2010).

49. Wen, L., et al. Whole-genome analysis of 5-hydroxymethylcytosine and 5-methylcytosine at base resolution in the human brain. Genome Biology 15, R49 (2014).

50. Elliott, H.R., et al. Differences in smoking associated DNA methylation patterns in South Asians and Europeans. Clin Epigenetics 6, 4 (2014).

51. Tsaprouni, L.G., et al. Cigarette smoking reduces DNA methylation levels at multiple genomic loci but the effect is partially reversible upon cessation. Epigenetics 9, 1382–1396 (2014).

52. Zeilinger, S., et al. Tobacco smoking leads to extensive genome-wide changes in DNA methylation. PLoS One 8, e63812 (2013).

53. Usui, N., et al. Foxp1 regulation of neonatal vocalizations via cortical development. Genes Dev 31, 2039–2055 (2017).

54. Dobin, A., et al. STAR: ultrafast universal RNA-seq aligner. Bioinformatics 29, 15–21 (2013).

55. Wang, L., Wang, S. & Li, W. RSeQC: quality control of RNA-seq experiments. Bioinformatics 28, 2184–2185 (2012).

56. Anders, S., Pyl, P.T. & Huber, W. HTSeq--a Python framework to work with high-throughput sequencing data. Bioinformatics 31, 166–169 (2015).

57. Leek, J.T., Johnson, W.E., Parker, H.S., Jaffe, A.E. & Storey, J.D. The sva package for removing batch effects and other unwanted variation in high-throughput experiments. Bioinformatics 28, 882–883 (2012).

58. Ritchie, M.E., et al. limma powers differential expression analyses for RNA-sequencing and microarray studies. Nucleic Acids Res 43, e47 (2015).

59. de Leeuw, C.A., Mooij, J.M., Heskes, T. & Posthuma, D. MAGMA: generalized gene-set analysis of GWAS data. PLoS Comput Biol 11, e1004219 (2015).

60. Davies, G., et al. Study of 300,486 individuals identifies 148 independent genetic loci influencing general cognitive function. Nat Commun 9, 2098 (2018).

61. Bipolar, D., Schizophrenia Working Group of the Psychiatric Genomics Consortium. Electronic address, d.r.v.e., Bipolar, D. & Schizophrenia Working Group of the Psychiatric Genomics, C. Genomic Dissection of Bipolar Disorder and Schizophrenia, Including 28 Subphenotypes. Cell 173, 1705–1715 e1716 (2018).

62. Jansen, I.E., et al. Genome-wide meta-analysis identifies new loci and functional pathways influencing Alzheimer’s disease risk. Nat Genet (2019).

63. Wray, N.R., et al. Genome-wide association analyses identify 44 risk variants and refine the genetic architecture of major depression. Nat Genet 50, 668–681 (2018).

64. Martin, N.W., et al. Educational attainment: a genome wide association study in 9538 Australians. PLoS One 6, e20128 (2011).

65. Savage, J.E., et al. Genome-wide association meta-analysis in 269,867 individuals identifies new genetic and functional links to intelligence. Nat Genet 50, 912–919 (2018).

66. Hoffmann, T.J., et al. A Large Multiethnic Genome-Wide Association Study of Adult Body Mass Index Identifies Novel Loci. Genetics 210, 499–515 (2018).

67. Schunkert, H., et al. Large-scale association analysis identifies 13 new susceptibility loci for coronary artery disease. Nat Genet 43, 333–338 (2011).

68. Morris, A.P., et al. Large-scale association analysis provides insights into the genetic architecture and pathophysiology of type 2 diabetes. Nat Genet 44, 981–990 (2012).

69. Estrada, K., et al. Genome-wide meta-analysis identifies 56 bone mineral density loci and reveals 14 loci associated with risk of fracture. Nat Genet 44, 491–501 (2012).

70. Boldog, E., et al. Transcriptomic and morphophysiological evidence for a specialized human cortical GABAergic cell type. Nat Neurosci 21, 1185–1195 (2018).

71. Satija, R., Farrell, J.A., Gennert, D., Schier, A.F. & Regev, A. Spatial reconstruction of single-cell gene expression data. Nat Biotechnol 33, 495–502 (2015).

72. Wang, X., Park, J., Susztak, K., Zhang, N.R. & Li, M. Bulk tissue cell type deconvolution with multi-subject single-cell expression reference. Nat Commun 10, 380 (2019).

73. GTEx Consortium. Human genomics. The Genotype-Tissue Expression (GTEx) pilot analysis: multitissue gene regulation in humans. Science 348, 648–660 (2015).

